# Improved methods for multi-trait fine mapping of pleiotropic risk loci

**DOI:** 10.1101/054684

**Authors:** Gleb Kichaev, Megan Roytman, Ruth Johnson, Eleazar Eskin, Sara Lindstroem, Peter Kraft, Bogdan Pasaniuc

## Abstract

Genome-wide association studies (GWAS) have identified thousands of regions in the genome that contain genetic variants that increase risk for complex traits and diseases. However, the variants uncovered in GWAS are typically not biologicaly causal, but rather, correlated to the true causal variant through linkage disequilibrium (LD). To discern the true causal variant(s), a variety of statistical fine-mapping methods have been proposed to prioritize variants for functional validation. In this work we introduce a new approach, fastPAINTOR, that leverages evidence across correlated traits, as well as functional annotation data, to improve fine-mapping accuracy at pleiotropic risk loci. To improve computational efficiency, we describe an new importance sampling scheme to perform model inference. First, we demonstrate in simulations that by leveraging functional annotation data, fastPAINTOR increases fine-mapping resolution relative to existing methods. Next, we show that jointly modeling pleiotropic risk regions improves fine-mapping resolution relative to standard single trait and pleiotropic fine mapping strategies. We report a reduction in the number of SNPs required for follow-up in order to capture 90% of the causal variants from 23 SNPs per locus using a single trait to 12 SNPs when fine-mapping two traits simultaneously. Finally, we analyze summary association data from a large-scale GWAS of lipids and show that these improvements are largely sustained in real data.

## Introduction

Genome-wide association studies (GWAS) have identified thousands of regions in the genome containing risk variants for complex traits and diseases [9, 29, 25, 31, 21]. However, the vast majority of the GWAS reported variants are not biologically causal, but rather, correlated to the true causal variants through linkage disequilibrium (LD) [30, 14, 16]. Fine mapping studies gather detailed genetic information within the loci that have been implicated in GWAS [23, 17, 32] and statistically dissect these regions to prioritize variants according to probability of causality. The top variants resulting from this procedure may become candidates for functional validation [6, 24].

Many statistical methods for fine-mapping have been developed for the prioritization of causal variants. Standard approaches range from a simple ranking of SNPs based on their p-values to more sophisticated LD-aware ranking algorithms that quantify probabilities for variants to be causal [14, 4, 1, 16]. Initial probabilistic methods have assumed a simple model in which only one variant per locus is biologically causal [22], with more recent methods extending the statistical frameworks to accommodate multiple casual variants at risk regions [14, 4, 16, 15]. Although modeling multiple causal variants drastically increases performance, particularly at loci with evidence of multiple signals of association, it also presents a combinatorially challenging problem in performing inference in the model. That is, the likelihood formulation contains a model space size exponential in the number of variants at a locus, which clearly cannot be enumerated over for even a modestly-sized locus. To account for this combinatorial explosion, initial methods approximated the full likelihood by restricting the maximum number of causal variants allowed at a risk locus to a small number [14, 4, 16, 15]. More recent works [1] further improved computational efficiency by sampling likely causal models using stochastic search, leveraging the intuition that most of the terms in the likelihood computation have near negligible contribution. The authors demonstrated that this achieves drastic reduction in runtime with comparable fine-mapping accuracy relative to enumerative methods [1]. However, this was done in the context of a single fine-mapping locus and did not integrate multiple sources of information.

Many GWAS loci are known to be implicated in multiple related traits – a phenomenon that is observed in many phenotypic classes. For example, breast cancer and mammographic density [19], high density lipoprotein (HDL) and low density lipoprotein (LDL) [9], or rheumatoid arthritis and irritable bowel disease [20, 25] are all pairs of traits that share overlapping GWAS signals. Combining association signals at these pleiotropic regions may strengthen the signal from the causal variants that are impacting both traits. A standard approach used when combining association information across multiple studies is fixed-effects metaanalysis, which assumes that causal variants across studies share the same effect sizes. The random-effects model does allow for effect size heterogeneity, but it is poorly-suited for situations in which the variant has opposite effect sizes in the various phenotypes [27]. For this reason, multivariate analyses that jointly analyze association data from multiple phenotypes and account for effect size heterogeneity are beneficial – particularly for related traits that have opposing phenotypic consequences such as HDL and LDL [9].

Considerable effort has been put forth into characterizing the chromatin landscape across the entire spectrum of human tissues [34, 8, 18]. Most recently, the Roadmap Epigenomics consortium interrogated 111 cell types, charting histone modifications, DNA accessibility, DNA methylation, and gene expression, to produce genome-wide maps of functional elements [18]. Previous works have demonstrated that principled integration of such data can aid fine-mapping performance in the context of single and multi-population fine-mapping studies [16, 15]. Since related traits have been shown to share an underlying genetic basis [2] that localizes within similar functional classes [11], it is plausible that functional annotation data can also augment cross-trait fine-mapping.

In this work we propose a unified framework to perform fast, integrative fine-mapping across multiple traits. We integrate the strength of association across multiple traits with functional annotation data to improve performance in the prioritization of causal variants. Our approach makes the assumption that the same variants at the risk loci impact both traits though with potentially distinct effect sizes. A key advantage of our approach is that it requires only summary association data for each trait, thus avoiding the restrictions that arise from the sharing of individual-level data. To balance computational efficiency and accuracy we propose an Importance Sampling technique that provides guarantees for convergence, while relaxing the assumption of the maximum number of causal variants allowed at each risk locus.

Through simulations we show that our integrative method delivers well-calibrated probabilities for SNPs to be causal and improves fine-mapping performance relative to current state-of-the-art strategies. To our knowledge, the only existing method that performs joint mapping for pleiotropy while incorporating functional annotation data is GPA [5]. We show that our approach provides superior accuracy to GPA, likely due to the explicit modeling of LD in our framework. We illustrate the benefit of our proposed methodologies by fine-mapping pleiotropic regions of lipid traits in a GWAS of over 180K individuals [9].

## Methods

### Overview

Here, we introduce statistical methods for fine-mapping of pleiotropic loci with functional annotation data (see Figure 1). We build upon previous works [16, 15, 14] that make use of a Multivariate Normal (MVN) distribution to jointly model association statistics at all SNPs at the locus. This not only allows for the possibility of multiple causal variants at any risk locus, but also avoids the need to access individual level genotype data as LD can be approximated using the appropriate population-matched reference panel [7]. We integrate relevant functional annotation data through a prior probability for SNPs to be causal. We introduce an Importance Sampling procedure to improve computational efficiency over methods that enumerate all possible models of causal configurations.

**Figure 1:**
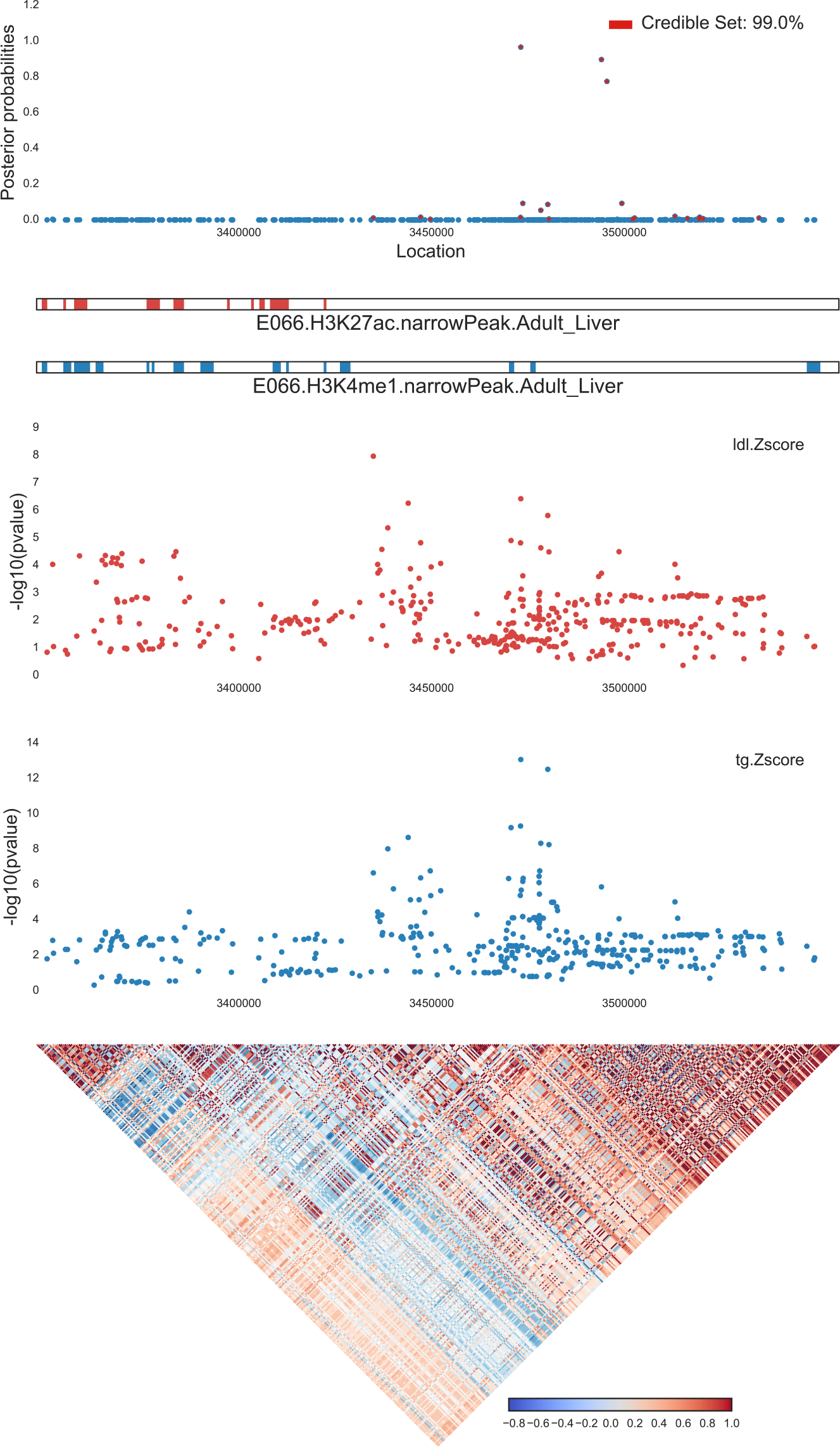
Example of input and output of fastPAINTOR at locus chr4:35Mb for LDL and TG. As input, fastPAINTOR receives an LD matrix, functional annotations, and multiple sets of Z-scores at the given locus. fastPAINTOR performs inference and outputs posterior probabilities for each SNP, indicating the likelihood that the SNP is causal across both traits.

### A statistical framework for fine-mapping

The standard approach to connect genotype to phenotype is through a linear model. For individual *i*, let *y_i_* be the trait value and ***g**_i_* be their vector of genotypes spanning *m* SNPs. The trait can be modeled as 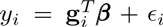, where 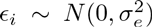 is random environmental noise. The vector, ***β***, represents the allelic effects whose entries will be non-zero only at the causal SNPs. Given *N* individuals with measured genotypes and trait values, the effect size at SNP *j* is typically estimated using standard linear regression as 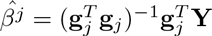. The strength of association is then quantified using the Wald statistic [3]:

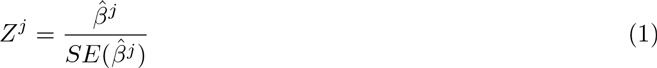

which asymptotically follows a normal distribution *Z ^j^* ~ 𝒩(λ*^j^*, 1) with mean

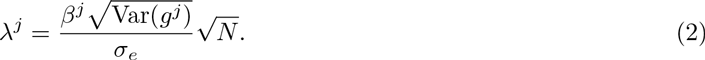

Here, λ*^j^*, is referred to as the Non-Centrality Parameter (NCP) and dictates of power of finding a significant association and, by extension, the power to distinguish causal from non-causal SNPs (i.e. β*^j^* ≠ 0 vs. β*^j^* = 0). When the *j*’th SNP is causal, the effect sizes are non-zero and therefore the association statistic (Z-score) corresponding to that SNP will be drawn from a non-central Normal distribution. However, LD (i.e. correlations between SNPs at each locus) will induce non-zero NCPs at non-causals variants through tagging. Therefore, neighboring non-causal SNPs will appear to be significantly associated to a trait indirectly through LD. Previous works [14, 16, 15] have shown that the NCPs at any SNP can be approximated from the NCPs at the causal SNPs:

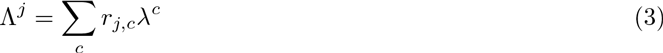

where *r_j,c_* denotes the Pearson correlation between SNP *j* and causal SNP *c*. If we collect all the pairwise correlations into a matrix, ∑, and let λ**_C_** be the vector of standardized effects sizes at the causal SNPs given by the indicator vector **C**, the entire set of regional summary statistics, **Z**, can be approximated by a Multivariate Normal distribution (MVN)) [14, 16]:

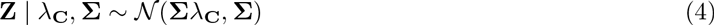

However, the causal effect sizes (λ**_C_**) are typically unknown apriori and must be either approximated [16, 15] or integrated out [14]. Leveraging the standard infinitesimal model [33], Hormorzdiari et al. [14] proposed to use a normal prior on the causal NCPs which, due to conjugacy, can be conveniently integrated analytically as follows:

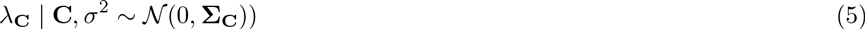

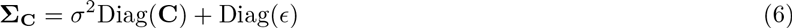

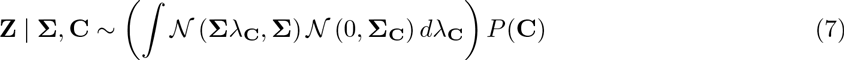

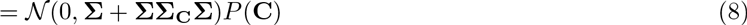

Here the prior probability of the causal set vector (*P*(**C**)) can be set to be uniform [22], hypergeometric [14], or can be estimated empirically using more sophisticated approaches that incorporate functional genomic data [16, 15]. Chen et al. [4] made the observation that the marginal likelihood in (eq. 8) is approximately proportional to a Bayes Factor comparing a causal and null model which depends on the Z-scores and LD only at the causal SNPs. This effectively reduces the computational burden from cubic in the number of SNPs to cubic in the number of causal variants considered at each likelihood evaluation. This not only improves efficiency, but also improves numerical stability since a much smaller matrix is inverted thus alleviating the need for stringent regularizations. In this work, we follow the Chen et al. implementation of the likelihood computations [4, 1].

### Fine-mapping pleiotropic loci

Next, we extend the framework to exploit pleiotropy across related traits. Given multiple phenotypic measurements across *T* traits, one can compute Z-scores for each trait independently. If a locus harbors a significant association for multiple traits, a reasonable assumption would be that the underlying causal variants driving this association are shared. It follows that the vectors of association statistics are conditionally independent given the causal variants (**C**), thus the joint distribution for all *T* sets of Z-scores decomposes into product:

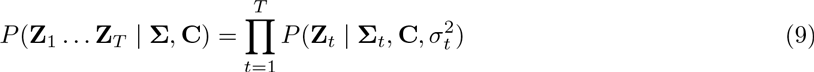

To simplify notation we hereafter refer to the collection of Z-scores at a fine-mapping locus as **Z**_*_ = {**Z**_1_ … **Z***_T_*}. We assume that all trait measurements have been performed in a single population and therefore assume that ∑*_t_*, = ∑ for all *t*. Importantly, we note that our formulation makes no assumptions on the coupling between effect sizes at causal SNPs across traits which allows for arbitrary levels of heterogeneity. Accommodating this effect size heterogeneity could be important for related traits that have opposing phenotypic consequences.

Under the assumption that causal variants are shared across pleiotropic loci, the marginal likelihood of the data can be written as a summation across all possible causal sets, 𝒞:

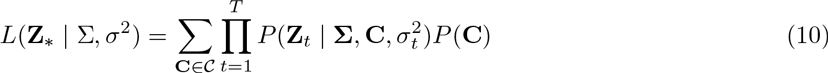

We can now use this to obtain the posterior probability of any causal set with a straightforward application of Bayes’ rule:

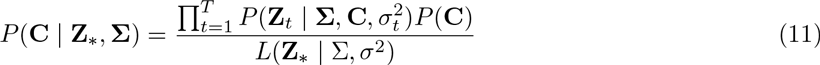

which can be marginalized to yield per-SNP posterior probabilities:

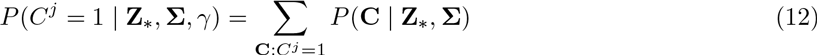

### Incorporating functional genomic data

To integrate functional annotation data within this framework, we use a logistic function to connect a SNP’s functional genomic context to its causal status as follows:

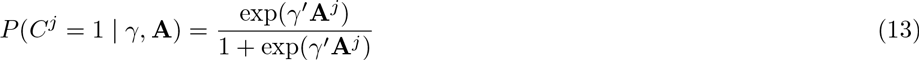

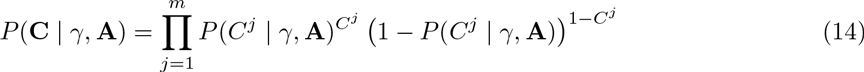

The vector **A***^j^* is the set of annotations corresponding to the *j*’th SNP and γ*_k_* is the prior-log odds that a SNP in annotation *k* is causal. We note that γ can be estimated directly from the data through an Empirical Bayes approach first described in Kichaev et al. [16]. However, this restricts functional enrichment estimation to only the fine-mapping loci under investigation. Alternatively, one could exploit potentially more powerful, genome-wide approaches such as stratified LD-score regression [11] that can infer global functional genomic enrichments using only summary data. Our framework is amenable to both approaches, and we allow the user to estimate γ from all the fine-mapping loci jointly using the EM algorithm proposed in [15] or supply it from external analyses.

### Model Inference via Importance Sampling

The marginal likelihood in (eq. 10) requires enumeration of *O*(2*^m^*) possible causal sets (𝒞). This rapidly becomes intractable as the number of SNPs grows large, and strategies for dealing with this computational bottleneck need to be considered. Earlier frameworks [16, 4, 15] avoided this problem by simply restricting the total number of potential casual variants to a small number (*k* << *m*), thus reducing the computational burden to *O*(*m^k^*). However, even in this reduced model space, enumerating over all possible combinations is inefficient as most causal configurations will contribute minimally to the overall likelihood of the data. Recent works have shown that sampling can circumvent brute-force enumeration by efficiently exploring likely causal configurations through stochastic search [1] – though this still requires pre-specifying a subjective prior that explicitly upper-bounds the maximum number of causal variants considered at the locus.

In this work, we make use of Importance Sampling, a variance reduction technique commonly used in Monte Carlo integration [12], to provide an efficient approximation of the marginal likelihood (eq. 10). Unlike other recently proposed sampling techniques, Importance Sampling comes with asymptotic convergence guarantees and allows us to drop the hard cutoff on the maximum number of potential causal variants considered. The summation given in (eq. 10) could naively be approximated by sampling directly from the prior and computing a simple Monte Carlo average:

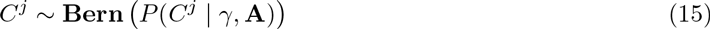

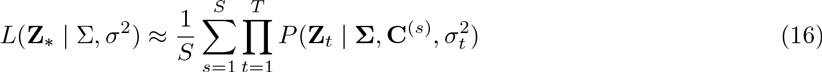

However, this is inefficient as highly probable causal sets in the posterior may not necessarily be reflected in the prior. To better guide the sampling of highly probable causal sets, we build off the intuition that SNPs with stronger associations are more likely to be casual than ones with weak associations. We can thus construct a discrete proposal distribution, *G*, to take this into account by simulating causal sets as independent Bernoulli draws with probabilities given by:

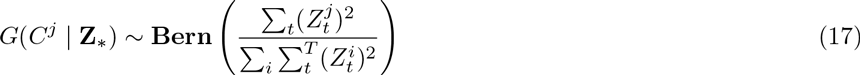

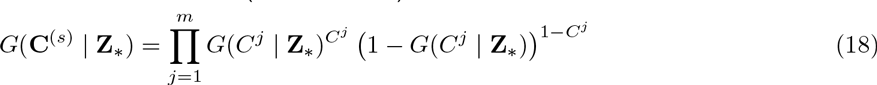

This proposal will favor selecting SNPs that have strong evidence of association in multiple traits. We can then compute importance weights and re-adjust the bias introduced by sampling from *G* as follows:

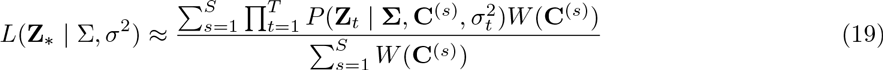

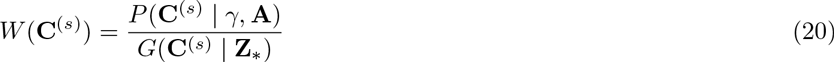

Which we can then use to approximate the per-SNP probabilities using the same *S* samples:

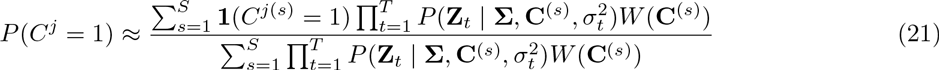

### Simulation Setup

To mimic real genotype data, we used HAPGEN2 [28] and the 1000 Genomes [7] European samples, to simulate 20,000 haplotypes for a number of randomly selected 25KB loci from chromosome 1. We filtered rare SNPs (MAF <0.01) and normalized genotypes to be mean-centered with unit variance. We overlapped our simulated regions with DNase Hypersensitivity (DHS) sites spanning 217 cell types and tissues [13]. Using these annotations, we drew causal status for each SNP according to the logistic model described previously, setting the DHS enrichment to 5.1 to reflect what was reported in [13]. Each locus harbored one causal variant in expectation, though the random assignment of causal status could yield zero or multiple casual variants for a given locus. In experiments that were done over 50 loci simultaneously, this typically resulted in an average of 18 loci with a single causal variant and 14 loci with multiple causals. Once we established the causal SNPs, we simulated phenotypes under a linear model such that for individual *i*, their phenotype value *Y_i_* was given by 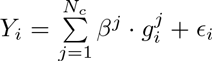, where *N_c_* is the number of causal variants, *β^j^* is the effect size of the *j*′th causal SNP, and 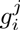 is number of copies of the risk allele *j* for individual *i*. We drew *ϵ_i_* for each individual from a normal distribution 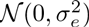, where 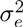 was given by the formula 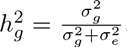, setting 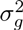 to the empirically observed genetic component.

We computed Z-scores for all the SNPs within causal loci by regressing the phenotype vector **Y** on each genotype vector **G***^j^* and then taking the Wald statistic. To simulate correlated traits, the effect sizes 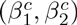 at the shared causal variants were drawn from an MVN distribution:

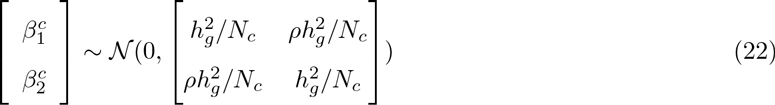

where *ρ* represents the desired genetic correlation. We chose a *ρ* of 0.4, consistent with typical correlations for lipids data reported in [2].

For computational efficiency, we also performed simulations in which the vectors of association statistics where drawn directly from an MVN distribution (eq. 4). In this scenario the NCP (λ**_c_**) was set to 5 at all causal SNPs.

### Existing methods

We compared our approach to several existing fine-mapping methods. For single-trait fine-mapping, we compared to FINEMAP and CAVIARBF [1, 4], two methods based on the CAVIAR[14] model that do not incorporate functional annotation data. We ran CAVIARBF v1.4 using the default settings, setting prior variance explained to be 0.05 and the maximum number of causal variants in the model to 3. After CAVIARBF computed Bayes factors for each SNP, we ran their model search algorithm, which outputs posterior probabilities based on Bayes factors. In this step, we set the prior probability of each SNP being causal to 1/*m*, where *m* is the number of variants in the locus. We ran the FINEMAP v1.1 software using default settings, allowing for 3 causal SNPs per locus with prior probabilities of (0.6, 0.3, 0.1) for 1, 2, and 3 causals respectively.

For multi-trait fine-mapping, we compared to GPA [5]. To our knowledge, GPA is the only other method that performs multi-trait fine-mapping while leveraging functional annotation data. As GPA requires p-values as input, we converted Z-scores from our simulations to p-values for each SNP. We provided GPA with the same DHS annotation data as we did for our approach. On multi-trait analyses, GPA outputs 4 posterior probabilities for each variant, indicating the probability that the SNP is causal for neither trait, Trait 1, Trait 2, or both traits. When evaluating accuracy, we considered the SNP to be deemed causal by GPA if it was implicated in both traits. In addition, we explored traditional meta-analysis techniques to combine information across traits by computing inverse variance fixed effects association statistics [10]. We then used these Z-scores in fine-mapping under the assumption of a single causal variant [22] as well as within our framework as a single trait.

### Empirical Lipids Data

We downloaded GWAS summary data across four blood lipids phenotypes: High Density Lipoprotein, Low Density Lipoprotein, and Triglycerides [9]. For each of the traits, we used Imp-G summary [26] to impute Z-scores up to the latest version (V3) of the 1000 Genomes European reference panel [7] yielding approximately 7.6 million SNPs per trait in total. We then compiled a list of 24 pleiotropic regions which we defined as a GWAS hit that was observed in least two traits of the three traits. For each of these regions, we centered a 250KB window around the lead SNP and overlapped these regions with two functional marks derived from the Roadmap Project: Liver H3K4me1 and Liver H3K27ac [18].

## Results

### Fast and reliable performance in single trait fine-mapping

We first sought to empirically assess how our sampling-based approach compared to fine-mapping methods CAVIARBF and FINEMAP. These previous approaches can model multiple causal variants, but were not designed to exploit pleiotropy. As such, in order to make the comparisons fair, we conducted our initial investigation in the context of a single trait. Furthermore, because these methods, as well as our proposed approach, are faster generalizations of the underlying CAVIAR model, we chose not to compare to CAVIAR nor PAINTOR, both of which would predictably have slower computational performance but similar accuracies.

We first assessed performance on the basis of CPU runtime. The number of samples that are drawn to approximate the posterior distribution is invariably connected to the resulting runtime for our method, fastPAINTOR. Therefore, we determined the number of samples required to yield approximately unbiased credible sets and find that one million samples was typically sufficient across a wide-range of locus sizes (Figure 2). We then compared to existing approaches and, not surprisingly, discover that methods that approximate the posterior model space through sampling vastly outperform methods that enumerate over all possible combinations (Figure 3). For example, both fastPAINTOR and FINEMAP scale favorably with the size of the locus, with average run times of (11.5s, 10.8s) per 25KB locus and (186s, 31s) per 250KB locus. The added computational overhead of fastPAINTOR is due to the fact that functional enrichments must be iteratively estimated using an EM-algorithm. If these estimates are supplied from external analyses, running fastPAINTOR* takes an average of 75s per 250KB locus to produce probabilities.

**Figure 2:**
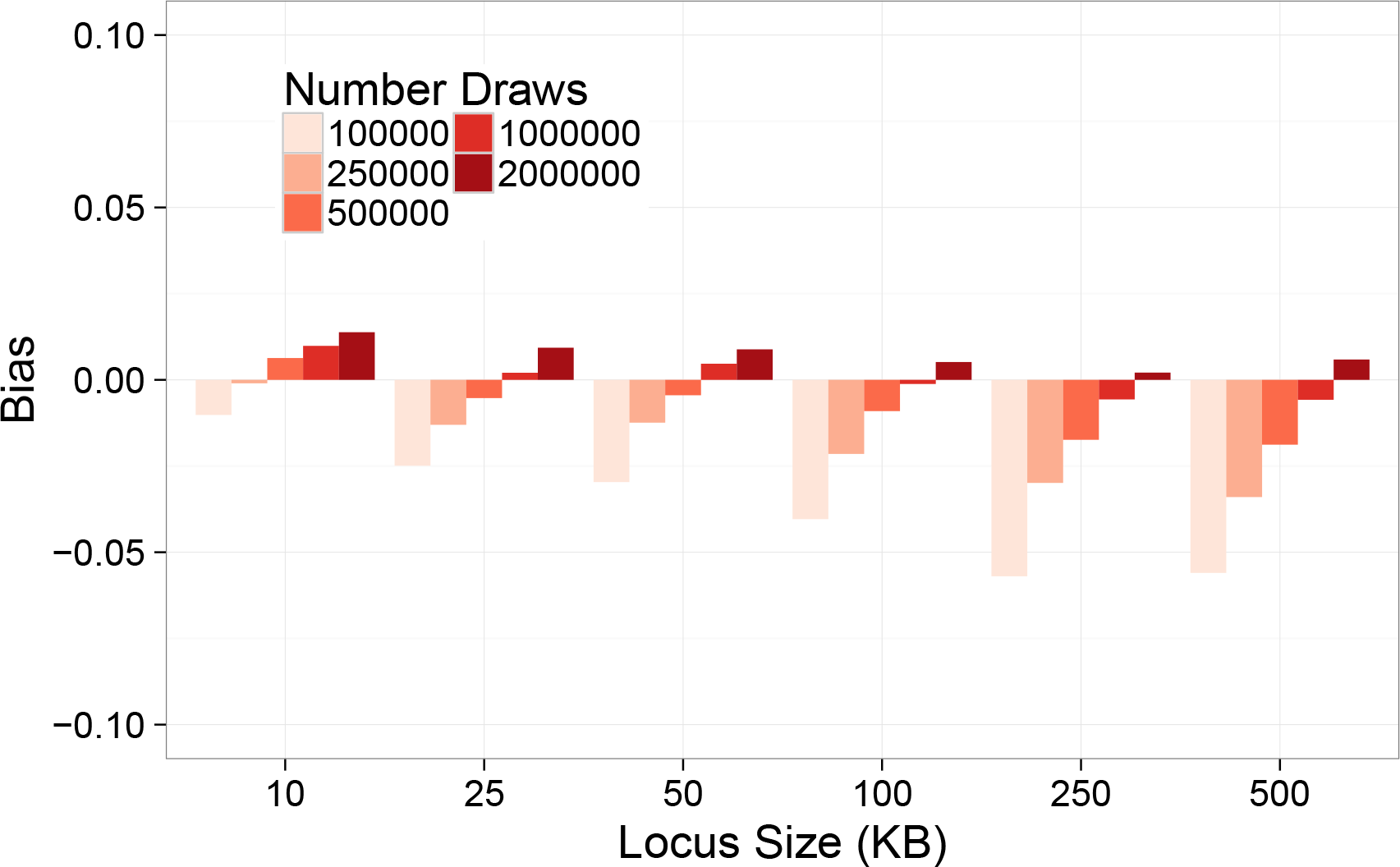
One million samples is sufficient to ensure approximately calibrated credible sets. We simulated variable sized regions by drawing from an MVN with reference LD given by the Europeans in the 1000 Genomes V3. We computed 95% credible sets for each simulated locus, and calculated the bias from defined as the difference between the proportion of simulated causal variants that were captured and the expected proportion (0.95). Here, negative bias represents a finding less causal variants than the credible set.

**Figure 3:**
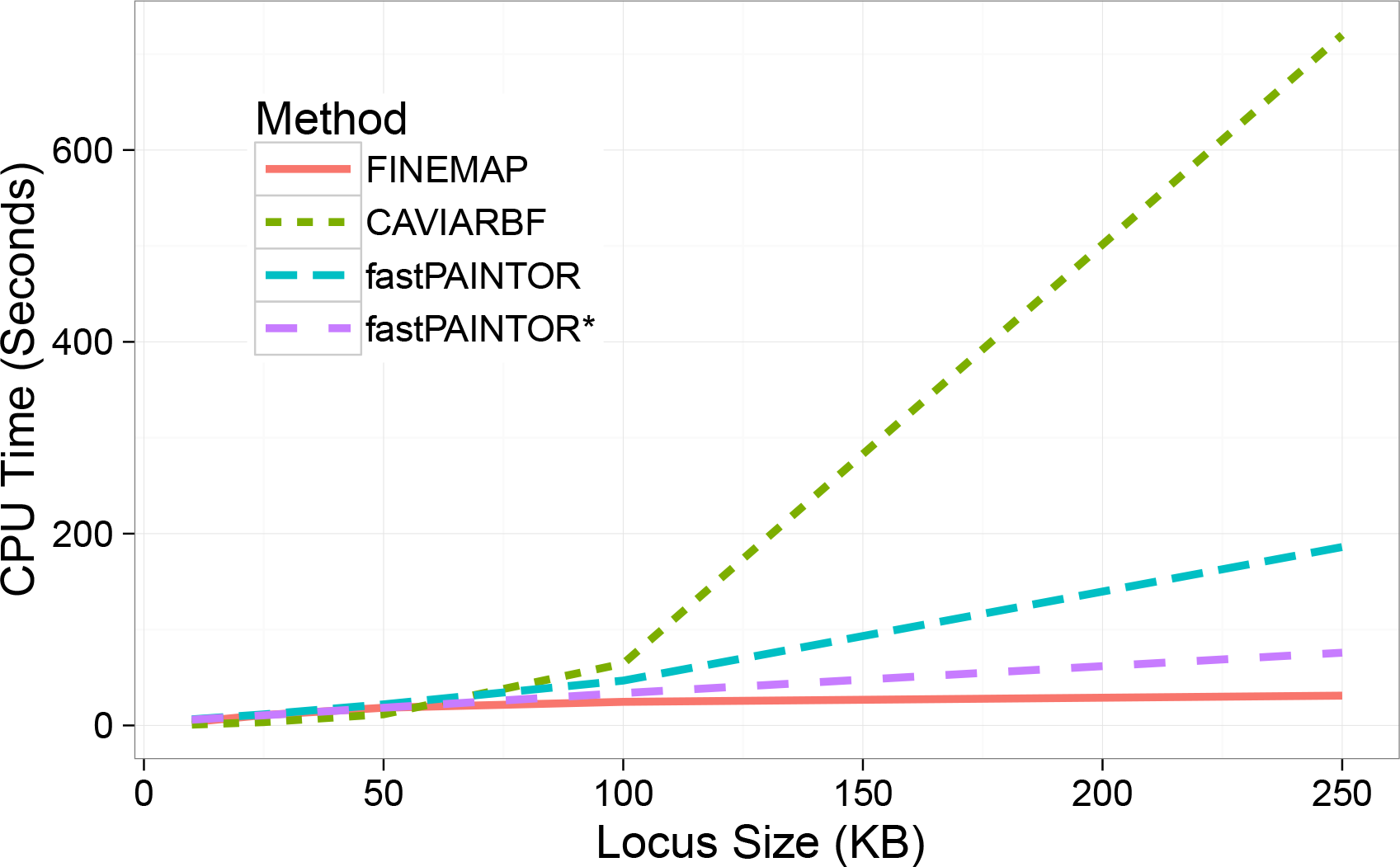
Importance sampling improves computational efficiency. Sampling approaches scale favorably with increasing number of SNPs being fine-mapped. We randomly selected 10 GWAS hits and centered increasingly large windows around them. For convenience, we simulated Z-scores by drawing from an MVN with reference LD given by the Europeans in the 1000 Genomes V3. Here, fastPAINTOR estimates functional enrichment empirically while fastPAINTOR* has it provided from external analyses.

We next evaluated the accuracy of these methods in resolving causal variants to ensure that our sampling approximation did not deflate performance. We simulated 100KB regions with various levels of DHS enrichment to reflect a wide diversity of potential functional genetic architectures. In general, we see that leveraging functional annotation data improves fine-mapping resolution relative to non-integrative approaches (Figure 4) – particularly as causal variants localize within smaller fractions of the genome (i.e. increasing enrichment). For example, the average rank of the causal SNPs was around 21.9 and 21.4 for CAVIARBF and FINEMAP across all functional genetics architectures. On the other hand, when causal variants are diffusely enriched within DHS, their average rank based on fastPAINTOR probabilities is 21.4 while strong functional enrichment yields an average rank of 15.0. Taken together, these results suggest that sampling-based, integrative methods are both scalable and achieve greater accuracy than current state-of-the-art methodologies.

**Figure 4:**
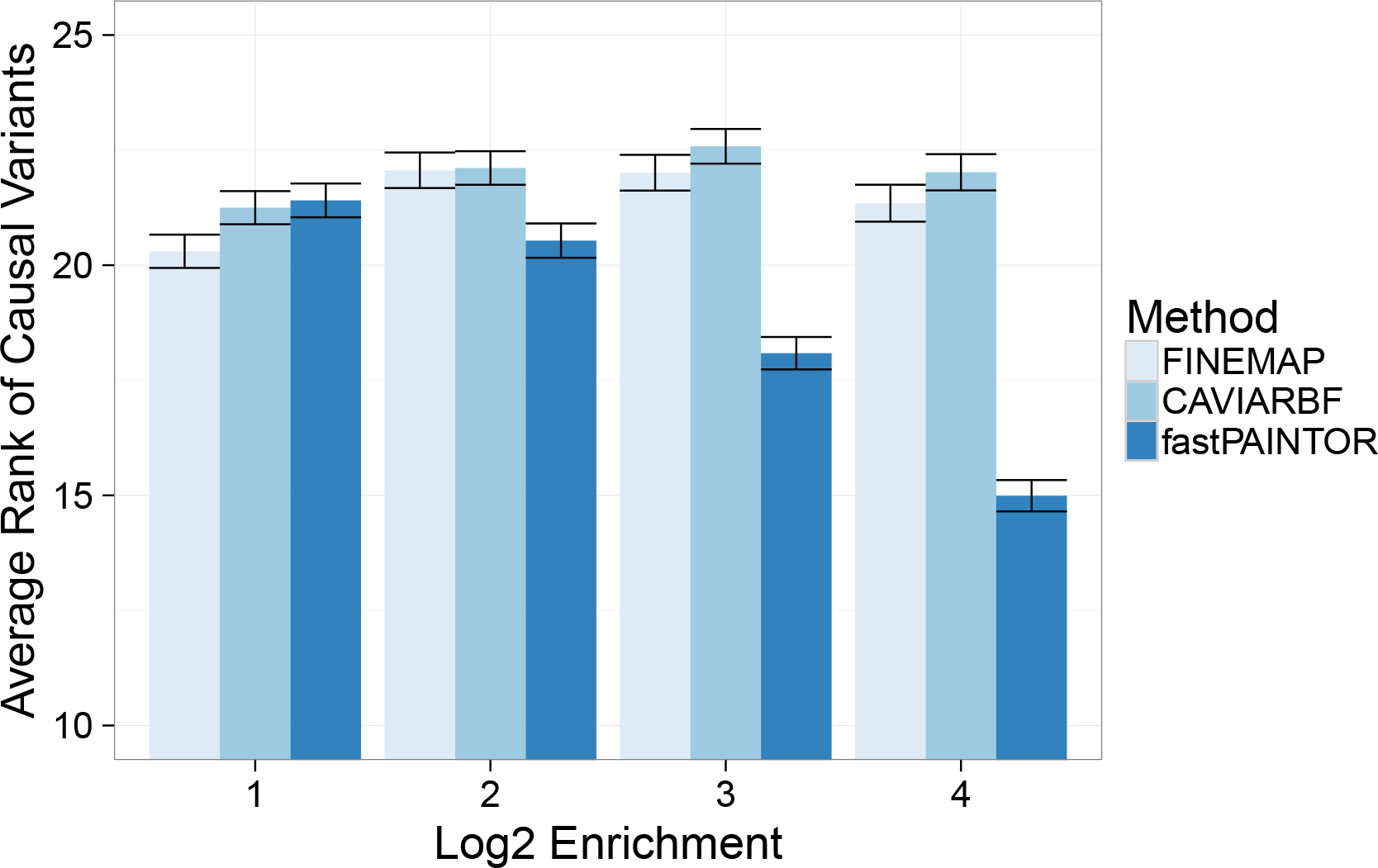
FastPAINTOR effectively leverages functional annotation data. We simulated fifty 100KB loci under various functional genetic architectures by drawing summary statistics directly from an MVN distribution. We applied all three methods using default settings and report the average ranks of the causal variants across all simulated loci.

### Multi-trait fine-mapping

Having established that our new computationally efficient approach compared favorably in standard fine-mapping scenarios, we next sought to investigate how leveraging information across related traits as well as functional annotation data affected fine-mapping performance. We simulated two traits for 10,000 individuals where the causal variants are shared between the traits but have heterogeneous effects (see Methods). We find that by borrowing information across related traits, we are able to improve fine-mapping performance with greater efficiency than just simply increasing sample size for any single trait (see Figure 5). In our multi-trait analysis with fastPAINTOR, we required (1.4, 12.4) SNPs per locus for follow-up in order to capture (50%, 90%) of the true causal variants, as compared with (1.9, 23.1) SNPs in a single-trait analysis. Intuitively, this is due to the fact that power to detect causal variants grows with the square root of the sample size, while growing linear with the allelic effects (see eq 2). Therefore, leveraging traits with multiple effect sizes will, on average, be more beneficial than simply increasing the sample size for one of the traits.

**Figure 5:**
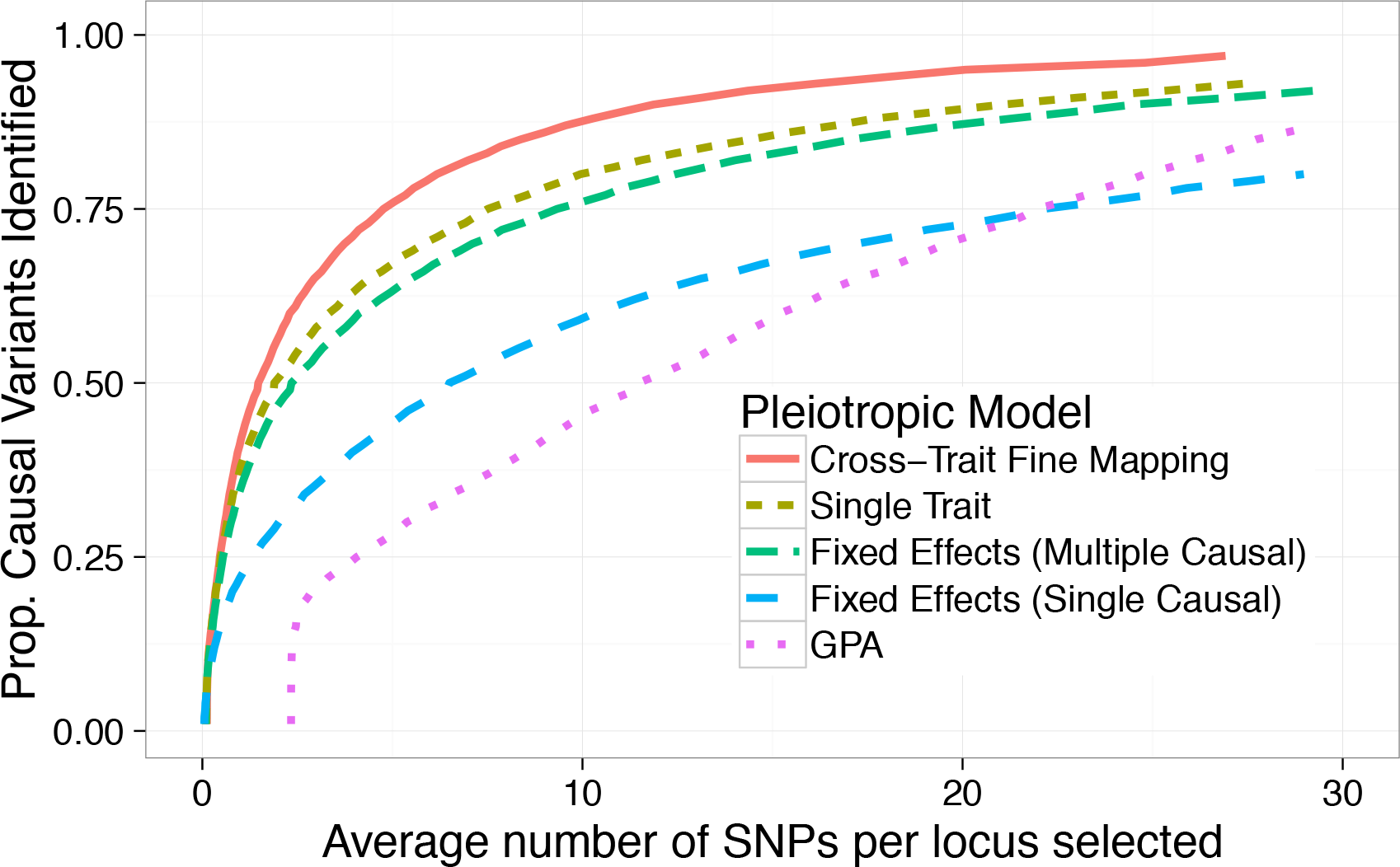
Integrative methods improve fine-mapping resolution in multiple traits. We simulated fifty 25KB loci for two traits with shared causal variants at each locus. We measure accuracy as the proportion of causal variants identified as we increase the size of our candidate SNP set.

We next explored principled strategies for assembling data spanning multiple traits. Our main comparator was GPA- a method specifically proposed to use pleiotropy and functional data to prioritize variants- as well as running fastPAINTOR with standard Fixed Effects (FE) meta-analysis. In general, our approach is more accurate and robust than previously proposed methods, requiring (1.4,12.4) SNPs per locus for follow-up in order to identify (50%, 90%) of the causal variants compared to (2.3,25.1) for fastPAINTOR with FE or (11.6,32.3) for GPA (Figure 5). One of the critical model assumptions of GPA is that SNPs are independent. Clearly, in the context of fine-mapping, this assumption is strongly violated which explains the sub-optimal performance. Alternatively, FE can be viewed as simply a weighted-average of the effect sizes. In the extreme, though not implausible, scenario where causal effects are going in opposite directions, FE will provide weak evidence that a SNP is causal.

Finally, we formulated our framework with the assumption that causal variants are shared across traits. This may not always hold in practice and we wanted to understand how our method responds to violations of this assumption. We performed simulations in which causal variants for the two traits were drawn independently leading to potentially distinct causal SNPs. We find that our joint fine-mapping method is robust to pleiotropic loci with differing causals, yielding relatively small mis-calibration of the credible sets on the order of 10% (see Table 1). We can thus conclude that our proposed framework that jointly models sets of association statistics, explicitly accounts for local correlation structure, and integrates functional data prioritizes variants robustly and accurately.

**Table 1:** The performance of fastPAINTOR is largely sustained when the assumption of shared causal variants across traits is violated. As compared with fine-mapping single traits independently, the reduction in the 95% credible set size is sustained while still capturing a large proportion of the causal variants. We define an 95% confidence set as the number of SNPs we need to select in order to accumulate 95% of the total posterior probability mass per locus.

| Method | Proportion of causals identified | SNPs selected (s.e.) |
| --- | --- | --- |
| Trait 1 | 0.96 | 46.01 (0.27) |
| Trait 2 | 0.96 | 45.54 (0.27) |
| Differing causals | 0.86 | 28.42 (0.22) |
| Same causals | 0.97 | 26.00 (0.17) |

### Multi-trait fine-mapping in lipids data

In order to demonstrate that the gains in our multi-trait fine-mapping approach are realized in real data, we analyzed summary association data from a large-scale GWAS of lipids [9]. High Density Lipoprotein (HDL), Low Density Lipoprotein (LDL), and Total Triglycerides (TG) are prototypical pleiotropic traits, sharing 24 GWAS hits for at least two. To showcase our pleiotropic fine-mapping framework, we obtained GWAS data over these traits spanning 180K individuals [9] and did integrative fine-mapping across putative pleiotropic regions. Functional annotation selection was guided by the genome-wide heritability-based functional enrichments reported in Finucane et al. [11]. The authors analyzed HDL, LDL, and TG and found that the H3K4me1 mark in liver tissue had the strongest enrichment of heritability across all three traits. Their result provides strong support for the key assumption that causal variants are shared across traits in our model. In addition to liver H3K4me1, we also used the liver H3K27ac mark, which displayed strong enrichment for multiple traits. In addition to a joint analysis, we applied our framework with and without functional data as well as on each trait independently. To quantify fine-mapping resolution we use 99% credible sets [22, 16] which are defined as the set of variants that aggregate to capture 99% of the posterior probability mass. Consistent with simulations, pleiotropic fine-mapping provided a reduction in the size of the credible set as compared with investigating individual traits alone (see Table 2). Additional functional data helps refine the signal, though only marginally, since exceedingly strong associations at these regions dominate the prior evidence. Moreover, we show that the 99% credible sets obtained from the cross-trait analysis contained 13 novel SNPs not found in any of the single-trait analyses alone (Figure 6). This suggests that, for some loci, leveraging association strength across related traits may increase our power to detect more weakly associated causal variants in the individual traits. In conclusion, these encouraging results illustrate that carefully merging related traits can improve the resolution of statistical fine-mapping.

**Figure 6:**
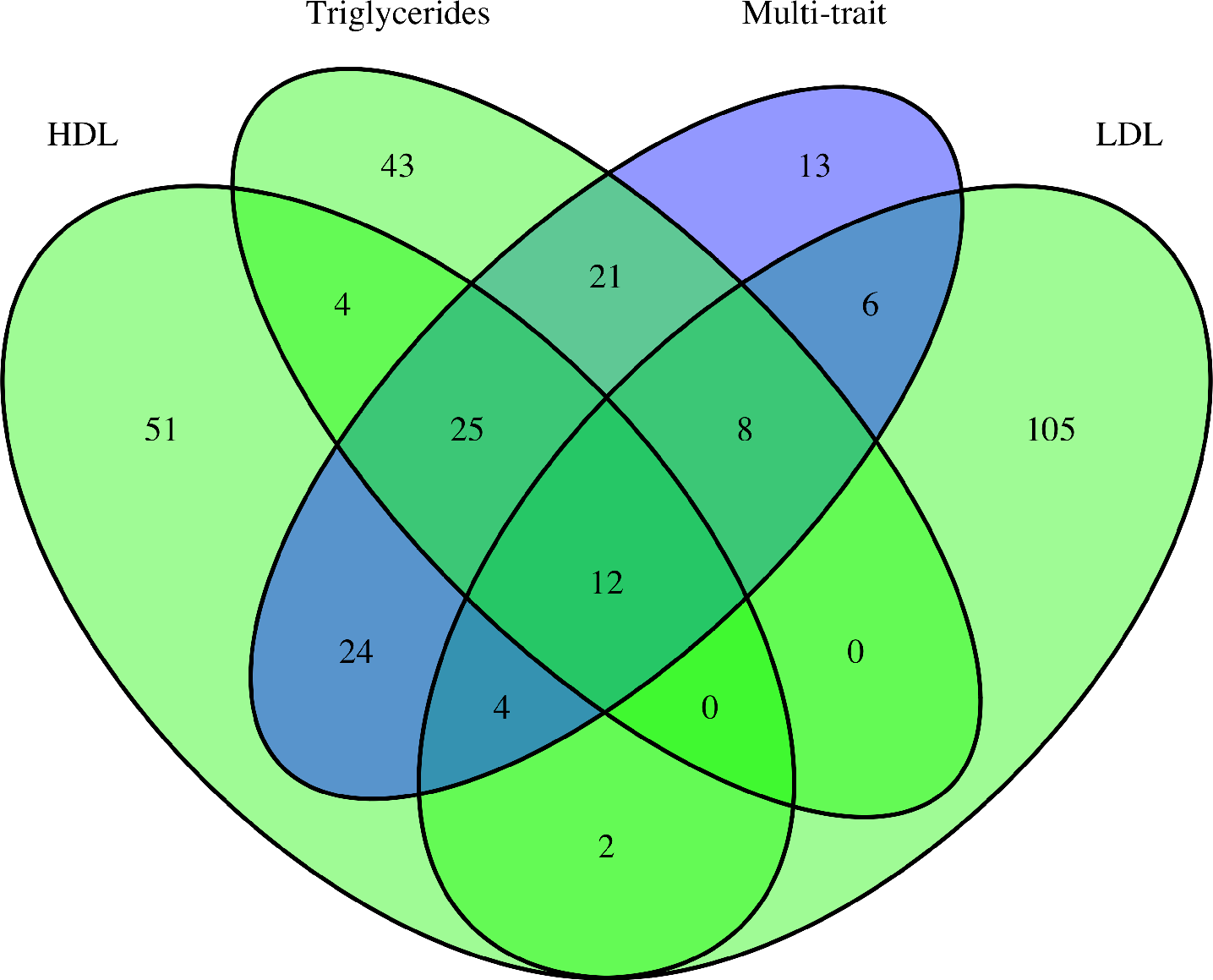
Cross-trait analysis proposes novel SNP sets. We obtained 99% credible sets for HDL, LDL, and TG analyses independently as well as for the joint analysis. We find that the credible sets from the cross-trait analysis contain 13 SNPs not found in any independent analysis.

**Table 2:** Pleiotropic fine-mapping is superior to single locus fine-mapping. Presented here are the mean number of SNPs that are in the 99% fine-mapping credible sets.

| Fine-mapping<br>Strategy | Annotations |  |
| --- | --- | --- |
|  | without | with |
| HDL | 4.83 | 5.08 |
| LDL | 14.25 | 11.42 |
| TG | 5.43 | 5.38 |
| Multi-trait | 4.71 | 4.71 |

## Discussion

In this work, we introduced a fast fine-mapping method that integrates several sources of genetic data to efficiently and accurately prioritize causal variants. Our Importance Sampling strategy dramatically reduces runtime due to its ability to efficiently sample high probability causal configurations, demonstrating that enumerating over complex model spaces is not necessary for integrative fine-mapping. We generalized this approach to leverage multiple traits simultaneously and demonstrated, both in simulations and real data, that this strategy can improve the ability to detect causal variants impacting both traits. As GWAS data accumulate and evidence for the abundance of pleiotropic risk loci mounts, there is a need for fine-mapping methods that can perform large-scale integrative analyses. Moreover, efforts by large consortia such as ENCODE will continue to provide genomic annotation data that will improve the accuracy of fine-mapping studies. A key advantage to our method is that it requires only summary association data, overcoming the issues that arise when sharing individual data that would otherwise limit sample sizes. In light of these developments, our proposed methodology will become increasingly applicable in the future.

We conclude by highlighting some caveats and limitations of our proposed framework. The power of our multi-trait fine-mapping framework hinges on the assumption that causal variants are shared at pleiotropic risk regions. While this notion is supported by the fact that related traits have shared functional genetic architectures [11], it is unknown whether this holds in general when doing fine-mapping. Reassuringly, we demonstrated in simulations that our framework is robust to this violation. Second, most large-scale GWAS have overlapping samples and the conditional independence assumption given in (eq. 9) may be violated. However, it is unclear whether this violation will bias the results dramatically if the underlying causal variants are shared across traits. Finally, while our Importance Sampling scheme does not explicitly upper-bound the number of causal variants at a fine-mapping regions, it favors exploring parsimonious models over complex ones. We therefore advocate that fine-mapping using our approach be undertaken where there is evidence of only moderate allelic heterogeneity.

